# The effect of silencing immunity related genes on longevity in the naturally occurring *Anopheles arabiensis* mosquito population of Southwest Ethiopia

**DOI:** 10.1101/361444

**Authors:** Serkadis Debalke, Tibebu Habtewold, Luc Duchateau, George K. Christophides

## Abstract

**Background:** In the fight against malaria, vector control remains the most important tool, butit is now severely constrained by the spread of insecticide or behavioral resistance by mosquito populations. Therefore, new vector control tools are warranted. Such novel tools include anti-mosquito vaccines or mosquito genetic modifications targeting the mosquito midgut homeostasis and reducing the mosquito lifespan beyond a stage they can transmit malaria.

**Methods:** We assessed the effect of RNA interference silencing of the midgut homeostasis regulators FN3D1, FN3D2, FN3D3, GPRGR9 and PGRPLC3 in populations of *Anopheles arabiensis* reared at nearly natural setting. We monitored the survival of gene-silenced mosquitoes and assessed the load of their midgut microbiota using flow cytometry. The effect of gene silencing was modeled by the Cox proportional hazards frailty model, and bacterial counts were first log transformed and then compared by a mixed model.

**Result:** Significantly higher mortality rates were observed for the *FN3D1* (Hazard ratio =1.64, P=0.004), *FN3D3* (HR=1.79, P<0.001) and *GPRGr9* silenced mosquitoes (HR=2.00, P<0.001) as compared to a control group injected with dsRNA against a non-related bacterial gene LacZ. The bacterial load ratios for all target gene silenced mosquitoes compared to control mosquitoes were above 1, with the highest value for *FN3D1* equal to 2.66 (95%CI: [0.94;7.57]) but no statistically significant difference could be demonstrated. Interestingly, there was a strong correlation (r=0.61) between the mortality hazard ratio and the bacterial count ratio of the gene-silenced mosquitoes. Increased mortality rates were reversed when the gene-silenced mosquitoes were treated with antibiotic mixtures suggesting that gut microbiota play a key role in the observed reduction of mosquito survival.

**Conclusion:** We demonstrate that interfering with the expression of *theFN3D1*, *FN3D3* or *GPRGr9* genes can cause a significant reduction of the longevity of *An. arabiensis* mosquitoes due to the disruption of the mosquito gut homeostasis.

## Authors summary

The mass distribution of insecticide-treated bed nets and the spraying of the inside of human dwellings with residual insecticides have significantly reduced malaria transmission throughout Africa in the last decade. However, both of the above tools exclusively target mosquitoes seeking blood meal and/or resting indoor, hence their ability to completely block malaria transmission is mostly incomplete. This is particularly problematic in regions such as Ethiopia where vector mosquito populations involve species that display behavioural plasticity with regard to feeding and resting site preferences such as *Anopheles arabiensis.* This reality stresses the need for developing novel vector control tools that can complement the existing interventions. In light of this, strategies involving transgenic mosquitoes and malaria transmission blocking vaccines, which interfere with the mosquitoes’ capacity to transmit malaria are considered. In this study, the authors examined whether suppressing the function of any of five genes expressed in the *An. arabiensis* midgut and involved in regulating bacterial homeostasis could reduce mosquito lifespan so much so that these mosquitoes do not live enough to transmit malaria from one person to another. They found that suppression of three of these genes could indeed substantially reduce the mosquito lifespan and that this reduction is dependent on the bacterial loads in the gut of these mosquitoes. This research was carried out in almost natural settings using mosquitoes collected from natural breeding sites, which reinforces the hypothesis that such interventions could be used to control malaria transmission.

## Background

Sub-Saharan Africa hosts some of the most efficient malaria vectors, namely the mosquitoes *An. gambiae, An. arabiensis* and *An. funestus*, and carries the heaviest malaria burden worldwide. Although slower compared to other disease endemic regions, a substantial reduction in malaria related cases (21%) and deaths (31%) have been recorded in the past decade [1]. This progress is largely attributable to the scale-up of vector control interventions, such along-lasting insecticide-treated nets (LLINs) or insecticide treated nets (ITNs) and indoor residual spraying (IRS), as well as to improved diagnostics for case ascertainment and effective treatments using artemisinin-based combination therapies [2].

LLINs/ITNs and IRS impact malaria transmission largely by reducing the daily survival rate of mosquitoes that are mostly active at night and display strong endophagic (seeking blood meal indoors) and endophilic (rest indoors following a blood meal) behavior [3-4]. [5] in their experimental hut trials demonstrated that despite having a similar sensitivity to insecticide *An. gambiae* was controlled more readily by LLINs than *A. arabiensis.*Structured experimental hut trials of ITNs and LLINs conducted in north-eastern Tanzania reported a consistently reduced mortality of *An. arabiensis* compared to *An. gambiae* and *An. Funestus* [5].Therefore, these tools have systematically reduced transmission by the endophagic/endophilic *An. gambiae* and *An. funestus* but not as much by *An. arabiensis* that exhibits significant exophagic (seeking blood meal outdoors) and exophilic (resting outdoors following a blood meal) behavior and thus remains partly resilient to the interventions[6-9]. In addition, field studies have reported evidence of behavioral adaptation (aka behavioral resistance) of *An. arabiensis* to the LLINs/ITNs byfeeding earlier and increasingly outdoors, and increasingly resting outdoors[7,10-11].

The aforementioned incompleteness of the efficacy current vector control tools has contributed to a marked shift in the dynamics of vector composition. Such shift is particular apparent in eastern African countries where these vectors coexist, *An. arabiensis* is observed to gradually replace *An. gambiae* and *An. funestus* [e.g. in Kenya [12-14] and Tanzania [7,17]. Although *An. arabiensis* is known to be a less efficient vector compared to *An. gambiae* and *An. funestus*, the inherent resilience of the mosquito to the LLINs/ITNs and IRS has been linked to reports of resurgence and stagnation, respectively, of malaria cases and deaths income African countries [e.g., 14-23], hence poses major threatens to the elimination goal. Therefore, there is an urgent need for additional vector control technologies targeting vectors that are resilient or have developed resistance to these tools.

Malaria transmission blocking vaccines and mosquito population replacement via genetic modification have recently become attractive alternatives to traditional vector control interventions [19-22]. Such interventions aim at rendering mosquitoes unable to be infected by and transmit malaria parasites [22-24] or reducing the mosquito lifespan beyond a stage that can transmit the malaria parasites to a new host [25-26]. Here, we screened *An. arabiensis* midgut genes as targets of mosquito life-shortening interventions that aim to generate mosquito populations with reduced lifespan. Our work was prompted by data showing that shortly after a blood meal the number of microbiota in the mosquito midgut increases drastically, to 100-1000 times [27-29], which in normal circumstances triggers immune reactions that soon reduces the microbiota number to the basal level [30-34]. We hypothesized that by compromising the immune system the mosquito can no longer control the microbiota in the gut, which would lead to a shorter lifespan. Previous studies have demonstrated that when genes encoding putative bacterial receptors such as PGRPLC, type III fibronectin domain proteins (FN3Ds including FN3D1, FN3D2, FN3D3) and the gustatory receptor GPRGr9 are silenced, gut homeostasis is disrupted in a laboratory colony of *An. gambiae* [31,35]. Here, we report a significant reduction in the longevity of naturally occurring *An. arabiensis* populations when some of the above genes are silenced by RNA interference. We conclude that interfering with the expression or disrupting the function of these genes using anti-mosquito vaccines or gene editing can lead to populations that cannot transmit malaria.

## Methods

### Experimental mosquitoes

Adult *An. arabiensis* mosquitoes were reared from larvae and pupae collected from natural breeding sites around Jimma, Southwest Ethiopia (7 40’ N, 36 50’ E). After visual inspection of the collection sites to check for the presence of mosquito aquatic stages, larvae and/or pupae were collected using a 350 ml capacity mosquito dipper following the standard larvae collection procedure [36-37]. The collected larvae and pupae were transported to an onsite mud-house where the pupae were separated from the larvae and transferred into a 10 ml beaker with water and kept in a mosquito cage (W24.5 × D24.5 × H24.5 cm, BugDourm-BD4F2222) until emergence. The remaining larvae were transferred in to a plastic tray containing rearing water obtained from their natural habitat and fed on yeast and fish food [36-37]. The larvae water was changed every 2 days and pupae were collected on daily basis and transferred into the adult cage. Emerged adult mosquitoes were maintained on 10% sugar solution. Two to three-day old adult females were transported to the experimental insectary at Jimma University for double stranded RNA (dsRNA) injection.

### Microclimate regulatory box

A special 2 m (length) × 1 m (width) and 0.75 m (height) box made from chipboard was constructed as a microclimate regulation structure to maintain experimental mosquitoes. The box was placed in a typical rural house and lined with 30 cm deep sawdust that was daily sprinkled with water to provide a condition similar to the natural resting mosquito habitat (S1. Figure). Experimental mosquitoes in paper cups were placed on the top of the sawdust, and the box was kept closed all the time with a lockable chipboard door. A window of 25 cm^2^was made at all four sides and the top cover of the box was covered with metal mesh to allow air conditioning. The box was placed on a 50 cm raised stand that was halfway dipped in water to provide protection from ant attacks. The box maintained a humidity of 60-70% RH and temperature of 25-28^0^C throughout the whole study period.

### Gene silencing

Total RNA was extracted from ten field-collected and laboratory reared *An. arabiensismosquitoes* using TRIzol (Invitrogen) and cleaned with Turbo DNase I (Ambion, UK). Complementary DNA (cDNA) was synthesized by reverse transcribing 1μg of the total RNA using Prime-Script^™^ 1^st^-strand cDNA Synthesis Kit (TaKaRa, UK). DsRNA for the five target genes was synthesized by PCR using specific gene primers tailed with the short T7 promoter sequence TAATACGACTCACTATAGG and the cDNA as a template. The targeted genes included*FN3D1* (AARA003032), *FN3D2* (AARA007751), *FN3D3* (AARA007751), *GPRGR9* (AARA003963) and*PGRPLC3* (AARA002982). DsRNA for the *LacZ* gene that served as a control was synthesized using as a template a plasmid containing the *LacZ* gene. The full primer sequences are shown in S1. table. DsRNA synthesis was carried out using the TranscriptAid T7 High Yield Transcription Kit (Thermoscientific, Lithuania). The dsRNA was then purified using the RNeasy Mini Kit (Qiagen, UK), following the manufacturer’s protocol. The concentration of dsRNA was determined spectrophotometrically by Nano drop 1000 spectrometer at 260nm and adjusted to 3μg/μl using ultra-pure water. Gel electrophoresis (1%TBE agarose) was performed on a subsample of the PCR products to confirm that products of the expected size were detected for each gene. Zero to two days old *An. arabiensis* mosquitoes were injected with 69 nl dsRNA specific to a target gene or the LacZ control gene following the RNA interference technique as described by [38].

Gene silencing efficiency was measured for each of the 5 silenced genes using qrtPCR. Quantification of transcript abundance was performed on cDNA synthesized from total RNA extracted from mosquitoes injected with dsRNA three days earlier and maintain on 10% sugar solution. Fast SYBR Green Master Mix used in the PCR reaction and the amplification was detected by a 7500 Fast Real-Time PCR system (Applied Bio Systems, UK). Each target gene was quantified in duplicate. The AgS7 gene was used as an internal control. Primer sequences are given in S 2. Table.

### Monitoring of mosquito survival

The dsRNA injected mosquitoes were put into their respectively labelled cups. For each gene between 20 and 30 mosquitoes were injected per replicate. The cups were kept in the microclimate regulatory box described above. The mosquitoes were offered a 10% sugar solution daily and a blood meal every fourth day by direct feeding on a goat. Survival was monitored daily for 25 days, starting 24 hours post injection. For each gene 6 replicates of mosquito injection were performed.

### Midgut microbiota analysis

For microbiota analysis, mosquitoes injected with the specified dsRNA were transferred into their respective labelled cups. For each gene between 20 and 30 mosquitoes were injected and were kept in a microclimate regulatory box. On day four post injection, the mosquitoes were allowed to feed on blood or kept on sugar meal. Twenty-four (24) hours post feeding four blood-fed and four sugar fed mosquitoes were sampled and their midguts were dissected. Individual midgets were homogenized in 100 μl 4% Paraformaldehyde (PFA) in Phosphate-buffered saline (PBS). The number of bacteria was quantified using flow cytometry on the five pooled midgut samples for a replicate [38]. A total of 3 replicates of dsRNA injections were carried out per gene.

### Antibiotic treatment

DsRNA-injected mosquitoes were placed in 6 different cups, each cup containing 20-30 mosquitoes. The cups were kept in the microclimate regulatory box. On the day of dsRNA injection, the mosquitoes were given a cotton ball soaked in antibiotic cocktail of Streptomycin and Norfloxacin, both a dose rate of 10 μg/mL in a 10% sugar solution. On day 4post injection, the mosquitoes were blood fed on a goat. The cotton balls were re-soaked with the antibiotic cocktail every fourth day for a period of 12 days (on days 4, 8and 12). A blood meal was also offered on the same days. The mosquito survival was monitored daily for 25 days starting from 24 hours post injection. For this experiment 5 replicates of dsRNA injection were performed per gene.

### Data Analysis

Data analysis was performed using the statistical software package R. For the gene silencing efficiency test, the relative expression of mRNA was calculated. First the threshold crossing values (Ct-values) of the samples were standardized using a standard curve obtained from the serial dilution curve and then normalized to the AgS7 gene transcript level. The relative expression of mRNA of a target gene was compared between mosquitoes for which the target gene was silenced and mosquitoes injected with control ds*LacZ* by a paired t-test using the replicate as a block factor. Gene silencing efficiency was expressed as the ratio of the relative expression of the target gene in the ds*LacZ* injected mosquitoes and the target gene silenced mosquitoes. Survival of the target gene and LacZds RNA-injected mosquitoes was depicted by Kaplan Meier survival curves. The effect of silencing the different target genes on survival was modeled by the Cox proportional hazards frailty model, with replicate used as frailty term [39]. The hazard ratio of a target gene over the control dsLacZ injection was used as summary statistic, together with the median time to death for the different gene silenced mosquitoes.

To investigate the effect of the antibiotics cocktail, the same Cox proportional hazards model was fitted using the hazard ratio again as summary statistic.

The bacterial counts were first log transformed and then compared by a mixed model with replicate as random effect, and the F-test was used to compare the silencing of the different target genes and the injection of control dsLacZ. The ratios of bacterial counts in the control dsLacZ injections and the target gene silencing were used as summary statistics.

## Results

### Effect of gene silencing on the survival

A significant reduction in expression level of mRNA for the specific gene silenced mosquitoes as compared to ds*LacZ*-injected mosquitoes was achieved for *FN3D1* (P=0.007), *FN3D2* (P=0.010), *FN3D3* (P=0.031) and *GPRGr9* (P=0.011) but not for *PGRPLC3* (P=0.145). The transcript levels of *FN3D1* (19%), *FN3D2* (20.5%), *FN3D3* (34.3%), GPRGr9(24%) and *PGRPLC3* (53%) were5, 4.9, 3, 4 and 2-fold lower compared to the ds*LacZ*-injected control group, respectively (Figure 1).

**Fig. 1:**
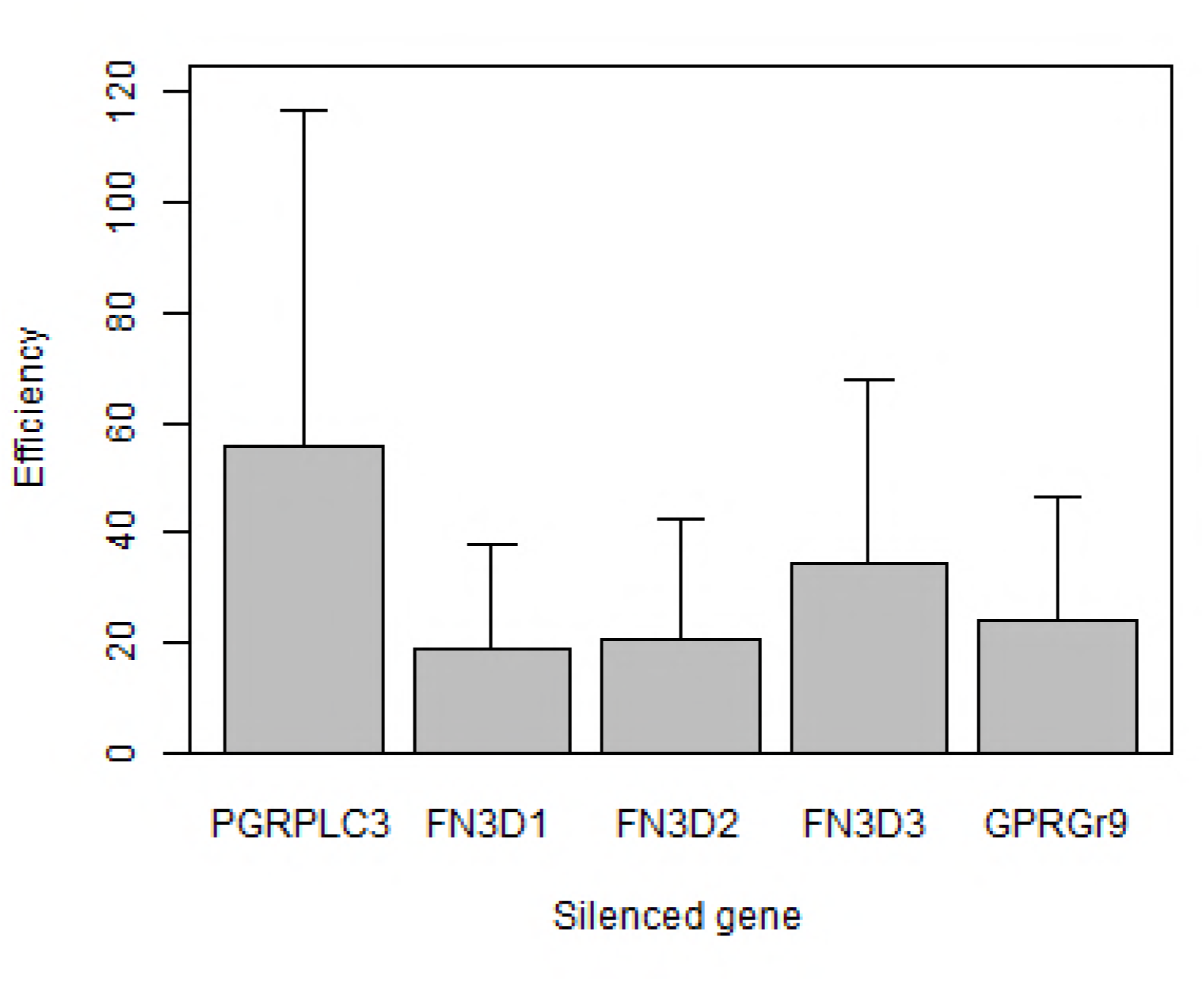
The average reduction (vertical line is standard error) in expression level of mRNA for the specific knocked down gene mosquitoes as compared to LacZ knocked down mosquitoes. For this experiment 3 replicates of dsRNA injections done.

The mosquito survival following silencing of the target genes compared to control ds*LacZ* injected mosquitoes is depicted as a function of time for the each of the above genes in Figure 2 Significantly higher mortality rates were observed for the *FN3D1* knocked down mosquitoes (Hazard ratio, HR=1.64), the *FN3D3* knocked down mosquitoes (HR=1.79) and the*GPRGr9*knocked down mosquitoes (HR=2.00) as compared to the control group, but not for *FN3D2* knocked down mosquitoes (HR=1.40),nor for *PGRPLC3* knocked down mosquitoes (HR=1.35) (Table 1).

**Fig. 2:**
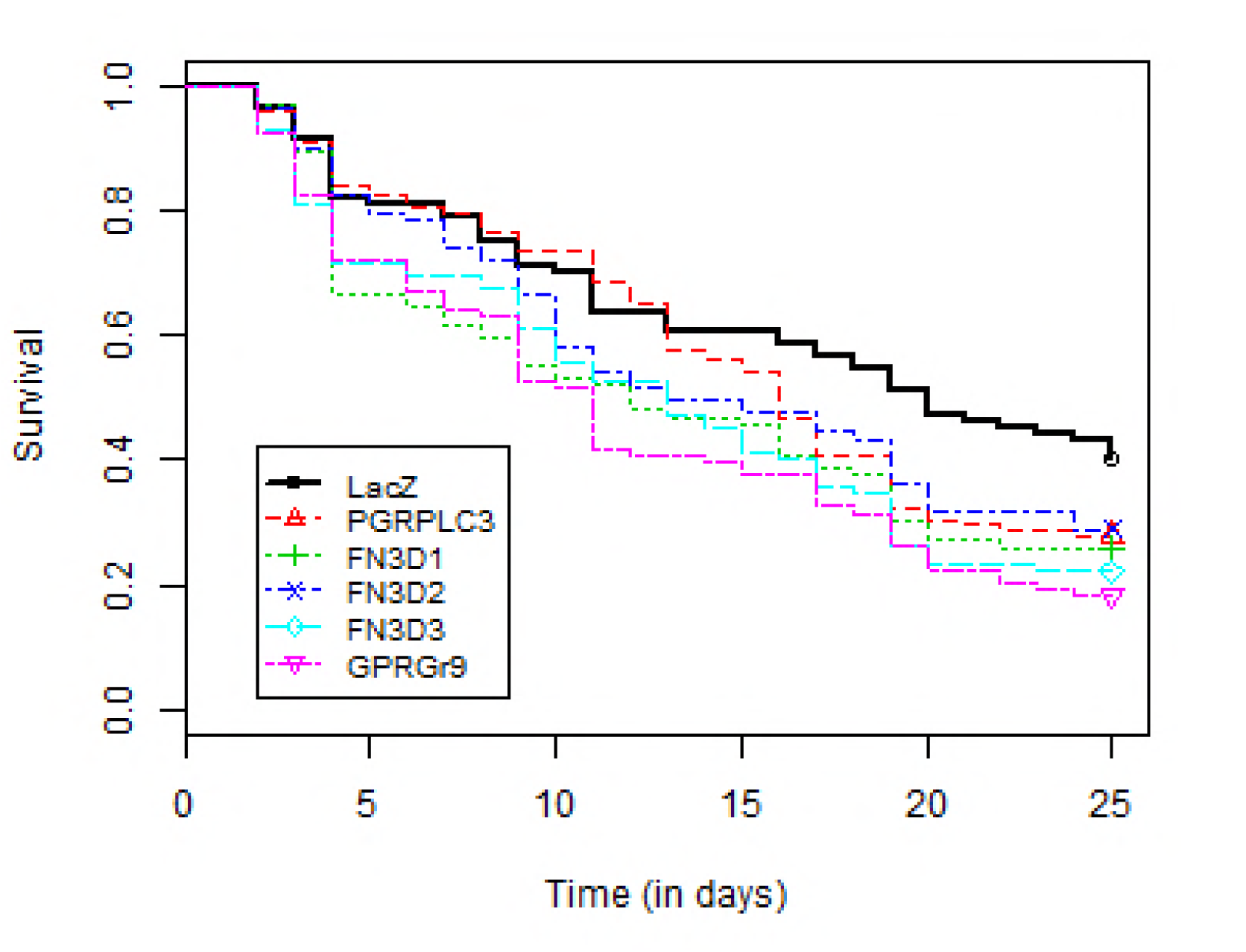
Kaplan Meier curves depicting the survival rate as a function of time for the 6 different gene silenced mosquitoes. The curves represented the survival rate of six independent biological replicates (n=6). In each replicate 20-30 mosquitoes were injected.

**Table 1.** Effect of gene silencing on mosquito survival. The second column presents the hazard ratio (HR) of dying between a particular silenced gene and ds*LacZ* control. The third column presents the hazard ratio of dying between a particular silenced gene and ds*LacZ* control when mosquitoes are treated in parallel with antibiotics.

| Genes | Hazard Ratio (95%CI; P-Value) |  |
| --- | --- | --- |
|  | Without antibiotics | With antibiotics |
| LacZ | 1.00 | 1.00 |
| FN3D1 | 1.64 ([1.17-2.30]; P=0.004) | 1.02 ([0.75-1.37]; P=0.91) |
| FN3D2 | 1.40 ([1.00-1.95]; P=0.050) | 1.17 ([0.86-1.58]; P=0.32) |
| FN3D3 | 1.79 ([1.28-2.50]; P<0.001) | 0.90 ([0.67-1.21]; P=0.50) |
| GPRGr9 | 2.00 ([1.45-2.76]; P<0.001) | 0.90 ([0.67-1.21]; P=0.47) |
| PGRPLC3 | 1.35 ([0.97-1.87]; P=0.072) | 0.89 ([0.66-1.21]; P=0.46) |

A marked reduction in average time to death was observed in mosquito groups where target genes were silenced compared to ds*LacZ*-injected controls. The median time to death was equal to 20 days (95% CI: [16;+∞]) for the control group and reduced to 12 days (95% CI: [9;17]) for the *FN3D1*silenced mosquitoes, to 13 days (95% CI: [10;17]) for the *FN3D3*silenced mosquitoes, to 11 days (95% CI: [9;15]) for the *GPRGr9* silenced mosquitoes, to 13 days (95% CI: [10;19]) for the *FN3D2*silencedmosquitoes and to 16 days (95% CI: [13;19]) for the *PGRPLC3*silencedmosquitoes. Gene silencing has no effect on the *Plasmodium falciparum* infectivity of the mosquitoes (S3. Figure) or mosquito sterility index (i.e. number of egg laid/female and egg hatchability), S4. table

### Effect of gene silencing on the midgut bacterial load

The global analysis demonstrated that there was no overall difference between the target gene silenced and the ds*LacZ*-injected mosquitoes with respect to bacterial load(P=0.363), but there was an overall effect of the feed source (P<0.001), i.e. blood versus sugar feeding, whereas the interaction between the factors was not significant (P=0.131). The ratio of bacteria load in blood-fed mosquitoes as compared to sugar fed mosquitoes was equal to 2.13 (95% CI: [1.49;3.06]).

No statistically significant differences in the bacterial load of the target gene silenced and the ds*LacZ*-injected mosquitoes were observed. The ratio of midgut bacterial count of target gene silenced to the ds*LacZ* injected mosquitoes is presented in Figure 3. The ratios for all target gene silenced mosquitoes compared to the control groups (LacZ injected mosquitoes) were above 1, with the highest value for *FN3D1* equal to 2.66 (95%CI: [0.94;7.57]) but not significantly different from 1 (P=0.085).

**Fig 3:**
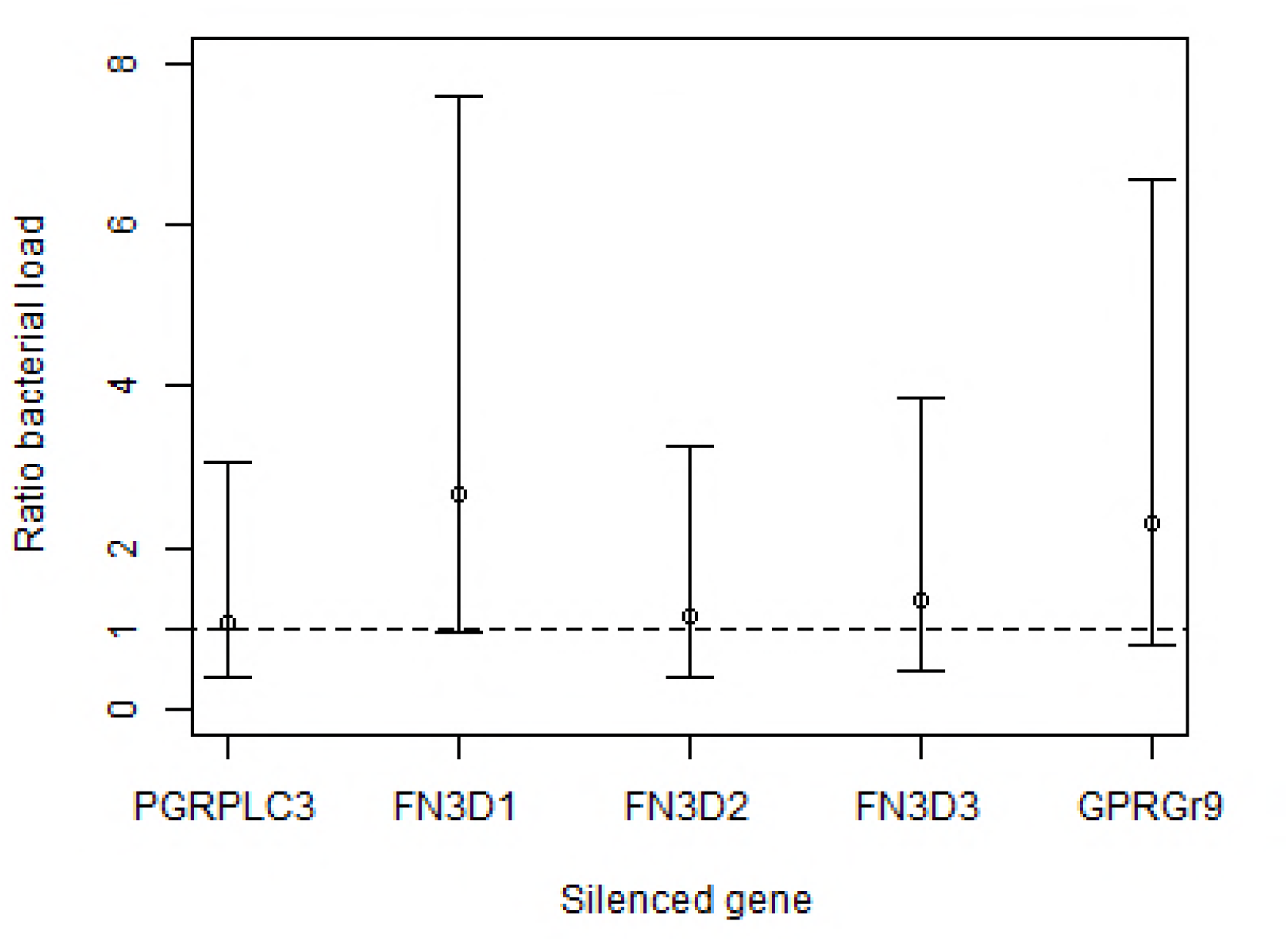
The ratio (95% confidence interval) of the bacterial load after 24 hours of micro-injection and blood meal in the 5 different target genes as compared to the LacZgene (n=3).

We further investigated whether there was a correlation between the mortality hazard ratio and the bacterial count ratio of the different silenced genes, i.e., whether a high mortality hazard ratio of a particular silenced gene corresponds with a high bacterial count ratio for that same gene. The relationship is presented in Figure 4, and it shows correlation (r=0.61), although it fails to be significantly different from 0(P=0.276). Never the less, treatment of mosquitoes with antibiotics cocktail eliminated the gene silencing effect on survival for all the 5 target genes. No statistically significant differences were noted in terms of mortality between the target genes silenced and the ds*LacZ* injected mosquitoes, with all hazard ratios being close to 1 (Table 1). This confirms that the reduced life span in gene silenced mosquitoes is related to disruption of gut microbiota homeostasis.

**Fig 4:**
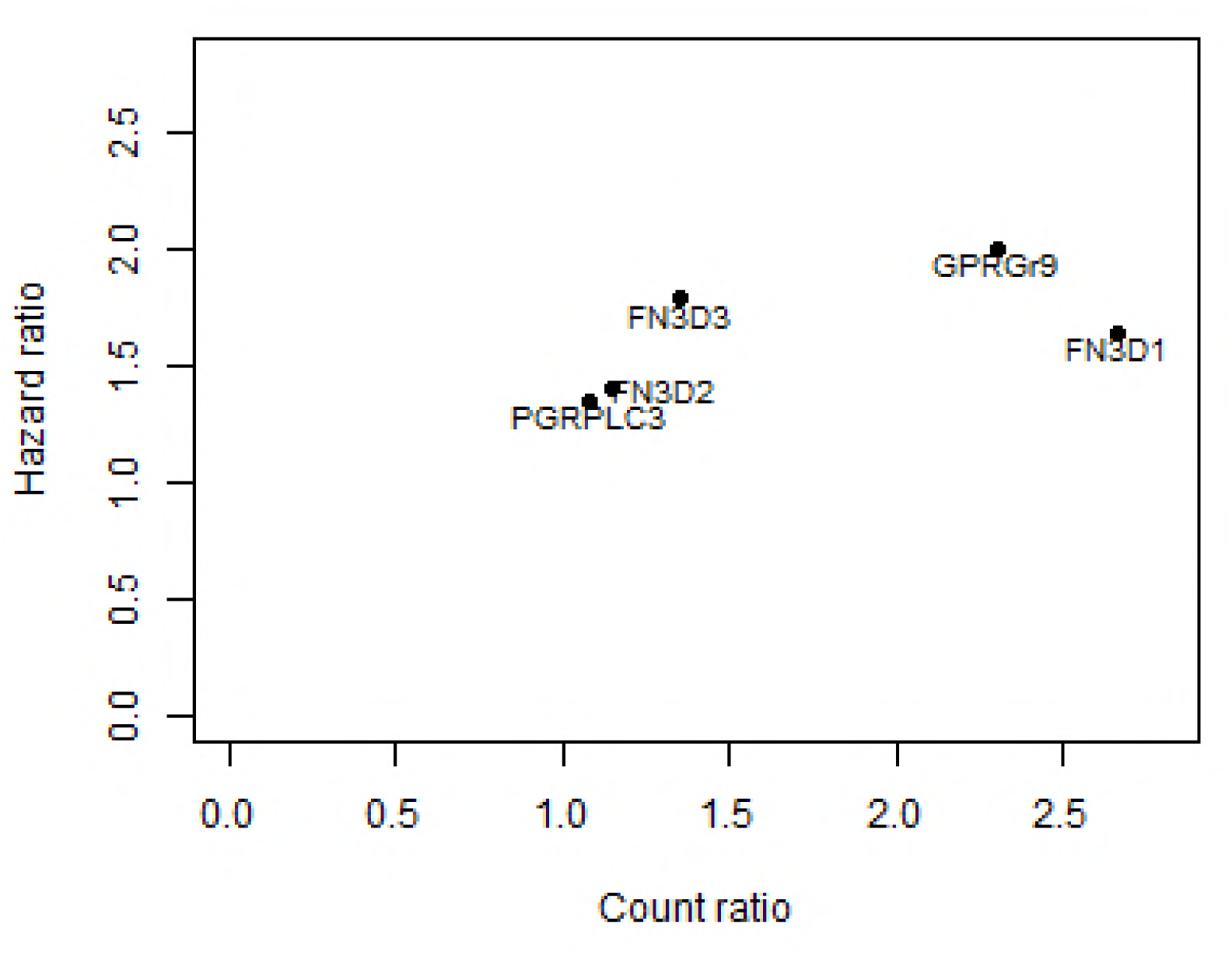
The relationship between the bacterial count ratio and the hazard ratio for the 5 different silenced genes.

## Discussion

New vector tools are required to complement the LLINs/IRS intervention to achieve the malaria elimination agenda in Africa. Of such novel approaches are releasing genetically modified mosquitoes and anti-mosquito vaccines delivered via alternative blood hosts. Genetic modification technologies that would aim to reduce the lifespan of mosquito populations can be an attractive option. A mosquito acquires the parasite when feeding on a person carrying *Plasmodium* gametocytes. Inside the mosquito, parasites go through a series of developmental stages and finally form sporozoites that reside in the salivary glands and are transmitted to a human host when the mosquito takes a blood meal. Depending on the parasite species and the ambient temperature, the required time for the parasite to complete its developmental stages inside the vector, known as the extrinsic incubation period (EIP), ranges from 10 to 21 days. Therefore, in order to be able to transmit the parasite, a female mosquito must survive longer than the EIP. Mathematical models indicate that the probability that a vector mosquito survives the duration of EIP is a critical factor in the vectorial capacity (C), i.e. the longer the mosquito lifespan the more efficient it is as a vector [40-41]. In nature, most mosquito species have lifespans that are too short to allow the parasite to complete its development, thus they are dead ends host. Even within a given species, most of the mosquitoes (90%) fail to survive this period mainly due to natural environmental and biological hazards, therefore any interventions that can reduce the mean mosquito survival period would offer a significant impact on the disease transmission [42-45].

In the present study, we have assessed five midgut-expressed genes *FN3D1*, *FN3D2*, *FN3D3*, *GPRGr9* and *PGRPLC* for their potential as lifespan limiting targets against a wild *An. arabiensis* population. Previous studies in*ane. gambiae* under laboratory settings have demonstrated that the above genes regulate midgut microbiota [31,35,38]. Unlike these studies, our work focused on the effect of those genes on the survival of field *An. arabiensis* populations. Our experimental mosquitoes were captured as larvae or pupae and maintained in an environment emulating the mosquito natural resting habitat. They were also allowed to blood feed on a goat every 4 day, fully reproducing their natural habits. We have demonstrated that in these almost natural conditions, reduction of*FN3D1*, *FN3D3* or*GPRGr9*expression significantly reduces the longevity of the *An. arabiensis* mosquitoes to an average of 12, 13and 11days, respectively, compared to a 20-day average longevity of control mosquitoes.

The observed effect on the mosquito survival can be attributed to the disruption of the mosquito midgut homeostasis. Although gene silencing did not affect the overall size of the natural midgut microbiota population to a statistically significant extent, two sources of evidence corroborate the role of microbiota in the observed reduction of mosquito survival. Firstly, a correlation (r=0.61) was observed between the mosquito mortality and gut microbiota load. Similarly, overall there was a higher microbiota count in the gene-silenced mosquitoes as compared to the control, although these differences were not statistically significant. Secondly, the observed effect of gene silencing (particularly of *FN3D1*, *FN3D3*, and *GPRGr9*) on the survival was reverted when mosquitoes were treated with an antibiotic cocktail to eliminate their gut microbiota. Our hypothesis that the reduced survival is due to the inability of mosquitoes to control their gut microbiota is also supported by other studies [46-47]. Furthermore, it has been previously demonstrated that *FN3Ds* and *GPRGr9*have a specific effect on microbiota of the *Enterobacteriaceae* family [35,38]. Hence, we consider that the observed survival phenotype is partly due to adverse alterations of the mosquito midgut ecosystem resulting in a disrupted or different state of homeostasis and physiology.

Our results suggest that interfering with the expression and/or function of these genes can reduce the mosquito lifespan which it turn will significantly impact malaria transmission due to reduced probability of the mosquito surviving the EIP. Such approach is supported by our observation in a separate experiment that the disruption of the expression of neither NF3D1 nor FN3D3 had a negative impact on the sterility index of female mosquitoes or their susceptibility to *P. falciparum* (data not shown) A permanent gene inactivation/knock-out can be achieved in various ways. A new and most promising tool is the recently developed CRISPR/Cas9-based genome editing methodology and gene-drive systems for *Anopheles* mosquitoes [24,48-49). Recently, [49] established a CRISPR/Cas9-induced somatic gene disruption technique in the *An. gambiae* mosquito. Such know-out line can be crossed with a germline-*Cas9* strain as described in [28,50] to generate germ-line gene-knockout line andthis line can be released to introgress the life-shortening trait into the wild malaria vector population and replace vector population [51]. A second approach can be through the immunization of the blood-providing host, whether human or other animals, against the target protein. Mosquito ingested antibodies would then neutralize the function of the protein leading to distorted midgut homeostasis and shortened lifespan. This approach is particularly attractive for zoophilic mosquitoes such as *An. arabiensis.* Mosquitoes that had an infectious blood meal (i.e. blood feed that is laden with Gametocytes) must take three-four additional blood meals before EIR as they normally blood feed every two-three day. This ensures repeated ingestion of anti-mosquito antibodies with consequential disruption of gut bacterial homeostasis to ultimately induce reduced lifespan. This approach has been successfully used for an anti-tick vaccine targeting the midgut antigens Bm86 and an anti-tick and anti-mosquito vaccine targeting the subolesin/akirin (SUB/AKR) antigens [52-55]. The functional model for the SUB/AKR vaccine involves the nuclear factor-kappa B (NF-kB) of vector insects to inhibit the Imd pathway, which is important for regulation of the gut microbiota [56-57].

## Conclusions

Here we demonstrate that interfering with the expression of the midgut proteinsFN3D1, FN3D3 or GPRGr9 significantly reduces the lifespan of *An. arabiensis* mosquitoesreared in nearly field conditions. The effect is caused by disruption of the mosquito midgut homeostasis through interference with the midgut microbiota, eventually hampering the mosquito immuno-metabolic functions. Therefore, these proteins can be good targets of mosquito life-shortening interventions, such as anti-mosquito vaccines or mosquito genetic modification, resulting in mosquitoes that can survive long enough to complete several gonotrophic cycles but not to transmit malaria parasites to a new host. Such technologies are expected to have a limited impact on the mosquito fitness thereby reducing selection pressure inducing resistance [58].

### Recommendation

We recommend further studies to dissect the impact of FN3D1, FN3D3 and GPRGr9 antibodies on the *An. arabiensis* longevity and midgut homeostasis.

## Abbreviations

cDNA: complementary DNA
Ct-values: threshold crossing values
DNA: Deoxyribonucleic acid
dsRNA: double stranded RNA
EIP: Extrinsic Incubation Period
FN3D1: Fibronectin type III domain-protein 1
FN3D2: Fibronectin type III domain-protein 2
FN3D3: Fibronectin type III domain-protein 3
GPRGR9: G protein coupled receptor protein 9
HR: Hazard Ratio
Imd: Immune deficiency
IRS: Indoor Residual Spraying
LLINs: Long Lasting Insecticide treated Nets
PGRPLC3: Peptidoglycan Recognition Protein LC3
PBS: Phosphate-buffered saline
PCR: Polymerase chain reaction
PFA: Paraformaldehyde
qrtPCR: Quantitative real-time PCR
RNA: Ribonucleic acid
TBV: Transmission Blocking Vaccines

## Ethical clearance

Ethical clearance was obtained from Institutional Review Board (IRB), Institute of Health, Jimma University.

## Consent for publication

Not applicable.

## Availability of data and material

Datasets are available from the corresponding author on reasonable request.

## Acknowledgements

We thank Professor Delenasaw Yewhalaw for kindly permiting us to use the facilities Tropical and Infectious Diseases Research Centre, Jimma University and Ethiopia, Mr Kassahun Ebba for logistic support.

## Supporting Information Legends

**S1. Fig:** Pictures of microclimate regulatory boxes.

**S2. Table:** list of primers used in the dsRNA synthesis and qPCR.

**S3. Fig:** *Plasmodium falciparum* oocyst intensity in gene silenced mosquitoes.

**S4: Table**: Fecundity and egg hatchability in gene silenced mosquitoes.

